# Impacts of a prolonged marine heatwave and chronic local human disturbance on juvenile coral assemblages

**DOI:** 10.1101/2024.02.23.581720

**Authors:** Kristina L. Tietjen, Nelson F. Perks, Niallan C. O’Brien, Julia K. Baum

## Abstract

Coral reefs are threatened by climate change and chronic local human disturbances. Although some laboratory studies have investigated the effects of combined stressors, such as nutrient enrichment and heat stress, on growth and survival of early life stage corals, *in situ* studies remain limited. To assess the influence of multiple stressors on juvenile corals, we quantified densities of corals ≤5 cm at 18 forereef sites with different exposure levels to underlying chronic local human disturbance before, during, and after the 2015-2016 El Niño. This marine heatwave caused prolonged heat stress and devastating losses of coral cover at our study site, Kiritimati, in the central equatorial Pacific Ocean. In total, we enumerated 7732 juvenile corals from 13 different families. Over 80% of corals were from four families: 70% from Agariciidae, Merulinidae, or Poritidae, which all have stress-tolerant life history strategies, and 11% from Acroporidae which has a competitive life-history strategy. Both local disturbance and heat stress were significantly negatively related to juvenile coral densities. Prior to the heatwave, juvenile densities were on average 72% lower at the most disturbed sites (7.2 ± 1.9 m^2^) compared to the least disturbed ones (15.3 ± 3.8 m^2^). Overall, juvenile corals had a lower bleaching prevalence and lower mortality during the heatwave when compared to their adult counterparts. Still, the heatwave resulted in the loss of half (50%) of all juvenile corals, with competitive and weedy life history strategy corals undergoing greater declines than stress-tolerant ones. Although juvenile coral densities increased slightly in the year following the heatwave, the effect was statistically non-significant. Our results highlight the influence of local chronic anthropogenic and climate change-driven marine heatwaves on juvenile coral densities.

## Introduction

Coral reefs are increasingly impacted by the effects of climate change, which are overlaid on chronic regional and local scale anthropogenic pressures [1,2]. Recent marine heatwaves, notably the 2015-2016 El Niño, have caused mass coral bleaching and extensive mortality [3–5]. The persistence of coral reefs is dependent on the recovery of corals, which can be driven by coral recruitment and the population dynamics of juvenile corals [6–8]. But while heatwave impacts have been documented extensively for adult corals, there has been comparatively less research on the effects of heat stress on juvenile corals. Previous studies have demonstrated that the demographics of juvenile corals can strongly influence recovery trajectories [6,7,9], with surviving juvenile corals rising through size classes, with corresponding increases in brood stock and reproductive output [10], to repair stock-recruitment relationships disrupted by the loss of adult colonies [10,11]. Studies of long-term reef recovery following the 1998 El Niño in the Indian Ocean (Scott Reef, Western Australia and the Maldives) found that juvenile corals stimulated reef recovery within 10-12 years of the event [10,12,13], and were likely especially critical on these isolated reefs that are reliant on self-seeding [10,14]. Overall, however, given their importance for recovery dynamics, and the lack of studies, there is a need to better understand the effects of climate change-amplified heat stress, and its interplay with local anthropogenic disturbance, on juvenile corals.

Although studies from several regions, including the Caribbean, Mediterranean, Japan, Thailand, and Australia [15–18], have shown that juvenile corals can have greater heat stress tolerance than their conspecific adults, the reasons underlying this difference remain unclear [15–17]. Mumby [15] hypothesized that this enhanced survival could be due to reduced irradiance levels due to their cryptic microhabitats or capacity for heterotrophic feeding to replace lost autotrophic nutrition during bleaching. Further research has suggested additional mechanisms based on properties such as being non-reproductive, which may allow for more energy invested into maintenance [18], and the relatively flat [19] and small colonies of juveniles which allow for faster elimination of toxic by-products by mass transfer [20]. In addition to exploring mechanisms for survival, a few studies have investigated the effects of heat stress on juvenile corals. Two experimental laboratory studies investigated the effects of short term heat stress and found that it resulted in sub-lethal stress and negative allometric growth scaling in *Porites* [21,22], whereas another study found increases in both growth and mortality in *Acropora* after over a month of elevated temperatures [23]. A comprehensive understanding of bleaching resilience mechanisms and heat stress effects in juvenile corals remains unresolved.

Juvenile survival through heat stress [7,18], and contributions to reef recovery through increases in coral cover [7], vary substantially amongst coral species and life history types. For example, a study by Doropoulos and colleagues [7] on the Great Barrier Reef, Australia found that when reef recovery is characterized by increases in coral cover, brooders may not considerably contribute to reef recovery due to their small colony sizes [7,24] despite their other ‘weedy’ life history characteristics (e.g., rapid generation times, opportunistically colonizing recently disturbed habitats) [24,25]. Rather, corals such as *Acropora*, that exhibit a ‘competitive’ life history strategy, grow large colonies, and excel at colonizing [24] can contribute considerably to coral cover [7], and thus play a major role in reef recovery. In comparison, massive corals that have a ‘stress-tolerant’ life history strategy often survive disturbance events [24], but do not contribute appreciably to increases in coral cover because of their slow growth rates [7].

Like adult corals, local factors, both natural and anthropogenic, can also affect the growth and survivorship of juvenile coral. A heat stress experiment investigated the influence of a simulated river plume and terrestrial runoff nutrient enrichment event on 4-month-old corals and found an antagonistic effect leading to reduced mortality rates [23]. This supports one of Mumby’s [15] hypotheses, as nutrients can be taken up by plankton communities which in turn are heterotrophically fed on by corals [reviewed in 26], potentially supplementing reduced autotrophic nutrition that is lost during bleaching [15]. However, on reefs in Barbados with human induced eutrophication and sedimentation, juvenile coral abundance was lower than on less eutrophic/low sediment reefs [27]. The effects of macroalgae may be the most well studied local factor on juvenile corals, and not unlike recruits or adult corals the effects are negative and result in slowed growth and decreased survivorship [21,28,29]. In turn, decreases in macroalgae levels have led to increased juvenile coral abundances [28,30]. Currently there is a lack of research on the impact of local factors have on the survival of juvenile corals which have important implications for reef recovery on chronically disturbed reefs.

Here, we aimed to fill a gap in the understanding of how multiple stressors affect the densities of different species of juvenile corals encompassing different life history strategies. To do so, we capitalized on a prolonged marine heatwave that occurred on Kiritimati (Christmas Island) during the 2015-2016 El Niño, which was overlaid on the atoll’s chronic disturbance gradient [31,32]. Kiritimati is geographically isolated, and thus is largely reliant on self-seeding for coral recruits [33,34]. We censused juvenile corals via video assays at 18 sites along the disturbance gradient before, during, and one year after the El Niño. This ecosystem-scale natural experiment allowed us to examine the impacts of prolonged heat stress on juvenile corals at sites exposed to different intensities of local human disturbance, and its effect on an isolated island’s initial reef recovery. We hypothesized that due to lower overall coral cover, decreased reef structural complexity and lower water quality [31] in areas with high human disturbance that, prior to the heatwave, these sites would have reduced juvenile coral densities. We also hypothesized that while elevated water temperatures would lead to a significant decline in juvenile corals: 1) bleaching and mortality would be reduced in juvenile corals compared to their adult counterparts due to their small morphology and microhabitats; and 2) mortality would vary amongst coral taxa, with a higher survival of stress-tolerant juvenile corals and increases in the year following the El Niño in weedy coral densities.

## Materials and Methods

### Study area and design

We quantified juvenile coral density by surveying their densities before, during, and after the 2015-2016 El Niño-induced heatwave, at eighteen shallow forereef (10-12 m isobath) sites spanning a gradient of local human disturbance on Kiritimati (Christmas Island, Republic of Kiribati). Kiritimati, a remote coral atoll in the central equatorial Pacific Ocean (01°52’N, 157°24’W), is the world’s largest atoll by land mass (388 km^2^; 150 km in perimeter; Fig 1). Sites were surveyed at three time points prior to impacts of the El Niño (two years before (July 2013; 16 sites), 1 year before (August 2014; 9 sites), and at the beginning (May 2015; 8 sites)), then resurveyed two months (early, July 2015; 13 sites) and 10 months (late, March 2016; 11 sites) into the heatwave, and again approximately 14 months after the end of the heatwave (1 year after; July 2017; 18 sites) (S1 Fig and S1 Table). Although not all sites were surveyed at each time point due to inclement weather conditions that prevented safe boat and/or diving access, the full set of eighteen sites were surveyed at least once before and once after the heatwave for direct comparability. The field research was conducted with permission from the Government of the Republic of Kiribati through permit numbers 008/13, 007/14, 001/16, 003/17.

**Fig 1.**
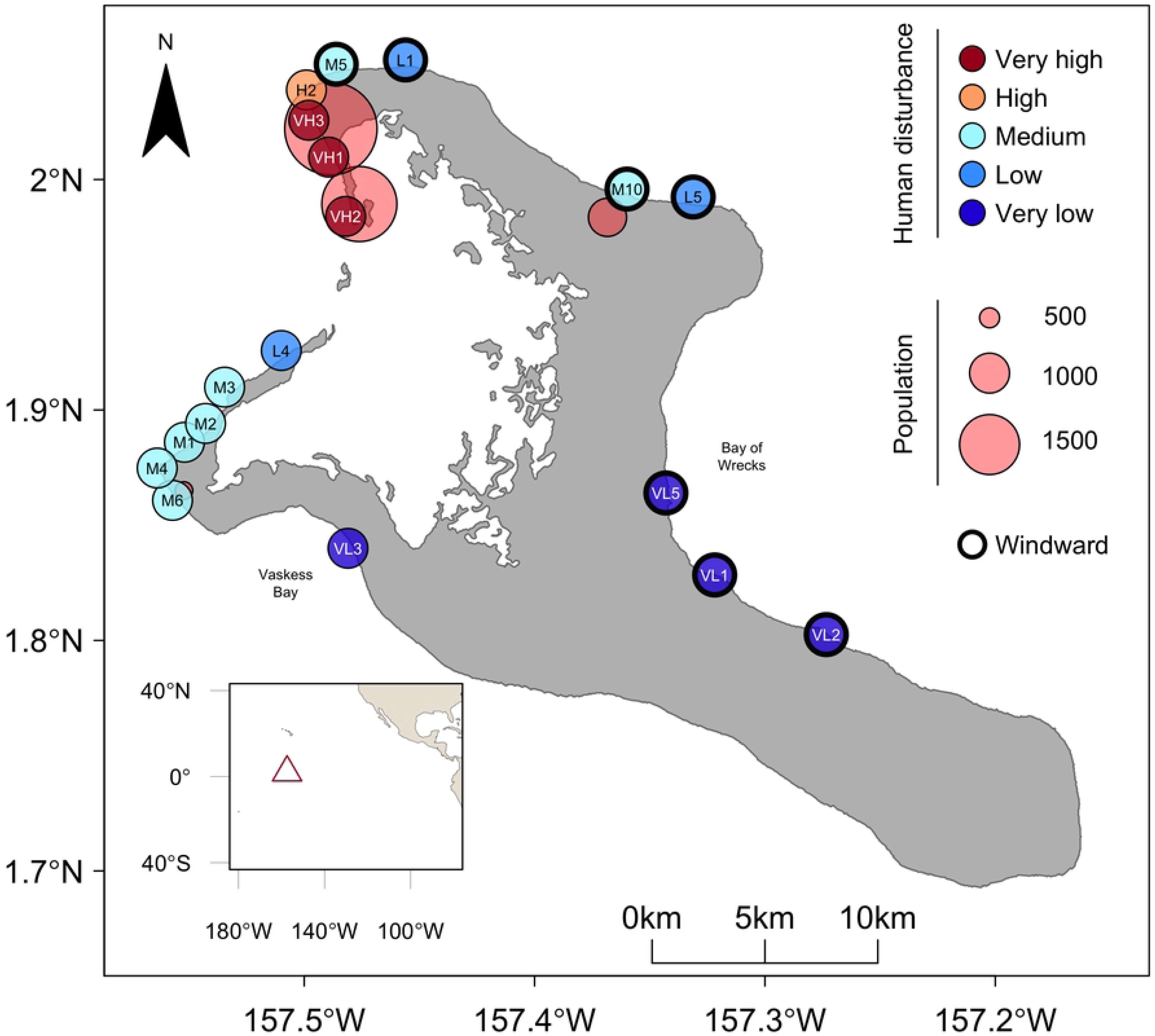
Map of shallow forereef study sites on Kiritimati (Christmas Island), categorized by level of chronic local human disturbance. Village populations (red circles) are represented by bubble size and windward sites are denoted by a thick black border around site. Inset shows Kiritimati’s location in the equatorial central Pacific Ocean (open triangle).

Chronic local human disturbance around the island was previously quantified [31,35] using a combined quantitative metric that incorporated both the human population within 2 km radius [36] of each site (as a proxy for localized impacts e.g. sewage inflow) [31,35] and fishing intensity data [32]. We modelled local human disturbance using this continuous metric, and for visualization used five local human disturbance categories (assigned previously from the continuous metric), which should be regarded as being relative to other sites rather than absolute levels of human disturbance [31]. Thermal stress, in degree heating weeks, was quantified from *in-situ* temperature data, collected from temperature loggers (SBE 56, Sea-Bird Scientific ± 0.001°C precision) deployed at monitored sites around the island, with at least one logger in each of the disturbance levels [further details in 37]. The heat stress from the 2015-2016 El Niño peaked at 27 Degree Heating Weeks (DHW) on Kiritimati, and temperatures were continuously elevated between June 2015 and April 2016 [37] with minimal differences in heat stress across sites (< 1.1 DHW) [31,37]. This prolonged heat stress resulted in an 89% loss of coral cover across the atoll [31].

In addition to local disturbance, oceanographic factors also vary around the atoll. We used site level net primary productivity (NPP; mg C m^-2^ day^-1^) data extracted from the Marine Socio-Environmental Covariates (MSEC) open source data product, derived from NOAA CoastWatch and calculated over a 2.5 acrmin grid (https://shiny.sesync.org/apps/msec/; [38]). In lieu of comprehensive site-level wave exposure data, we defined site-level exposures based on the dominate wind direction (southeasterly; [39]), with sheltered sites on the west side of the island grouped as leeward, and exposed sites on the north and east coasts as windward, following [31,35,40].

### Juvenile coral survey

At each site, we surveyed juvenile corals along two 25 m transects. Up to ten 1 m^2^ gridded quadrats were set at predetermined random points along the transects and filmed following the protocol in Mumby and colleagues [41], with the modification of 10 cm swaths instead of 20 cm (n = 732 videos total, S1 Table and S2 Fig); more than ten quadrats were surveyed at site VH1 to account for the high prevalence of sand at that site. Videos were randomly analyzed by one of three trained individuals (KLT, NFP, NCOB), who identified each coral to the lowest possible taxonomic unit and measured its 2D horizontal size (i.e., widest width) using the software Tracker (physlets.org/tracker/; [42]; Fig 2); KLT also checked identifications to ensure consistency amongst observers. A juvenile coral was distinguished as being a new colony—rather than a fragment of a colony that experienced partial mortality—by inconsistencies with the surrounding area (i.e., dead skeleton). We defined juvenile corals as those with ≤5 cm maximum width. While this size classification is somewhat arbitrary in regards to maturity of corals, it is a commonly used size classification in juvenile and young coral studies [e.g., 7,18,45,46]. Corals that were not fully inside a quadrat were removed from the dataset. When available, life history traits for identified corals were assigned as in Baum and colleagues [31], based upon the Coral Trait Database (https://coraltraits.org; [24]) to classify species found in the database, and extracted life histories for other species based on the family (S2 Table). Three coral taxa (n = 42 individuals total) could not be assigned to a life history strategy and thus were removed from statistical analysis. In addition, corals with unresolved identifications or life history assignments (e.g., when coral could only be identified to family and a life history could not be assigned to the family due to multiple strategies within the family) were removed from statistical analysis (n = 317; 4.1% of the data). Generalists (n = 5) and soft coral (n = 38) were excluded from our statistical analysis due to low densities.

**Fig 2.**
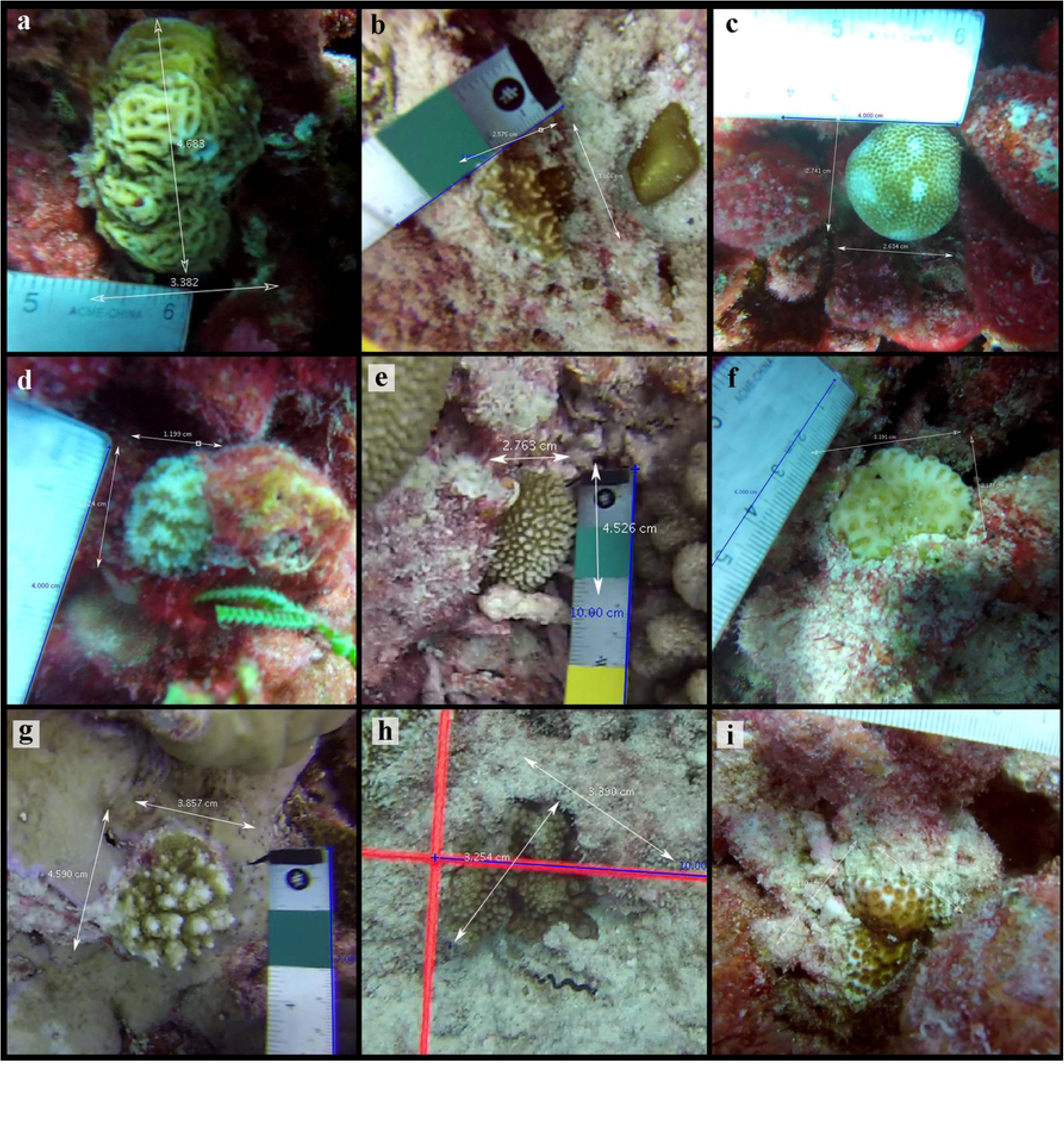
Representative photos of surveyed juvenile corals on Kiritimati throughout the study. The top three most common corals were **(a)** *Leptoseris mycetoseroides*, **(b)** *Pavona varians*, and **(c)** *Porites lobata*. Panels d-f (d = *Platygyra* spp., e = *Hydnophora microconos*, f = *Goniastrea stelligera*) are examples of other stress-tolerant corals. Panels g (*Acropora* spp.) and h (*Pocillopora* spp.) are examples of corals with competitive life histories, and panel i (*Leptastrea purpurea*) has a weedy life history strategy.

### Statistical analyses

We first examined how juvenile coral densities varied around the atoll prior to the heatwave, by fitting generalized linear mixed-effects models (GLMMs) to the data for the three pre-heatwave timepoints, using a negative binomial error structure to accommodate overdispersion in the data. We included an interaction between local human disturbance (continuous) and life history strategy, along with reef exposure (leeward, windward) and NPP, as fixed effects, while site was modelled as a random effect. Local human disturbance was modelled as a polynomial (quadratic) relationship to allow for the possibility of a non-linear relationship, although the quadratic was ultimately reduced to a linear due to AIC model selection (detailed below). We also included an offset to account for differences in the total number of quadrats sampled at each site.

We then tested for the influence of the heatwave on juvenile coral density by adding to the above models ‘heat stress’ as an additional fixed effect (four levels: ‘before’ heatwave = 2013, 2014, and May 2015; ‘early’ = July 2015, ‘late’ = March 2016; and ‘after’ = July 2017), and a two-way interaction between local disturbance (modelled as a quadratic) and heatwave period to examine if the heatwave impact was modulated by local disturbance. We conducted two sensitivity analyses to test if the heatwave effect was dependent on the exact sites sampled in the various expeditions. The first sensitivity analysis included only the 9 sites that were sampled in all four periods and the second included the 10 sites sampled in both the before and in the late heat stress periods (S4 Table and S1 Fig). Finally, we investigated if the impacts of the heatwave varied with coral life history strategy, by fitting separate ‘heat stress’ models for each life history type.

All statistical analyses were conducted in R v.4.0.3 [45]. GLMMs were tested using the package *glmmADMB* [46,47]. Prior to analysis, all continuous input variables were standardized to a mean of zero and a standard deviation of 0.5 using the ‘rescale’ function in the *arm* package [48]. AIC was used in model selection and diagnostic graphs plotting residuals using the *DHARMa* package [49] were analyzed for each model presented. For each model type, we present the top model according to AIC, accounting for model parsimony by selecting the model with fewer fixed effects when there were two top models within 2 AIC of one another.

## Results

In total, we enumerated 7732 juvenile corals (n = 7320 hard; n = 38 soft; n = 374 unidentifiable) from 732 census quadrat videos. Juvenile corals had an overall mean density of 10.7 m^2^ (± 0.74 SE) and a mean width of 2.28 cm (± 0.01 SE; S3 Fig). We identified 45 juvenile coral species (or genera when species was not possible) from 13 different families (S5 Table), the most common of which were *Leptoseris mycetoseroides* (2.1 ± 0.08 m^-2^), *Pavona varians* (1.3 ± 0.16 m^-2^), and *Porites lobata* (1.0 ± 0.10 m^-2^) (Figs 2 and 3). Over 80% of enumerated corals belonged to four families: Agariciidae (n = 2732), Merulinidae (n = 1874), Acroporidae (n = 873), and Poritidae (n = 774) (S5 Table). Stress-tolerant corals were the most common at every time point and accounted for almost three-quarters (73%) of the corals overall (n = 5661) (Figs 3 and 4). Competitive corals were the next most common (n = 1065 corals; 13.8%), followed by weedy (n = 550, 7.1%), and generalists (n=5) (Fig 3).

**Fig 3.**
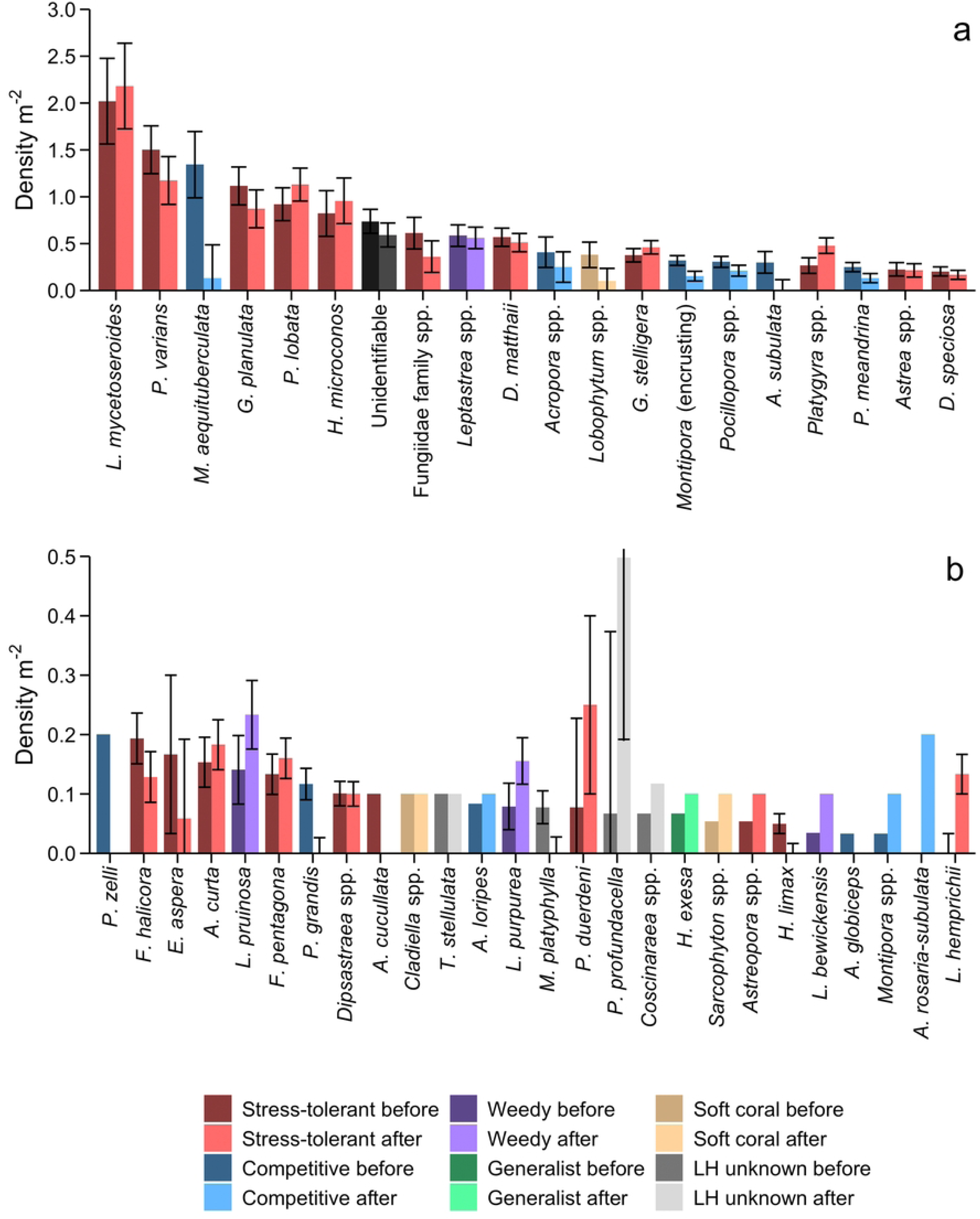
Density (± SE) of 45 juvenile (≤5 cm) coral species (or genus) before (2013, 2014, and 2015b) and after (2017) the marine heatwave. Plots **(a)** 20 most common and **(b)** rarer corals are colored by life history and heatwave period. Error bar for *P. profundacella* (in panel b) after the marine heatwave extends to 0.8042 m^-2^. Y-axis scales vary between plots.

**Fig 4.**
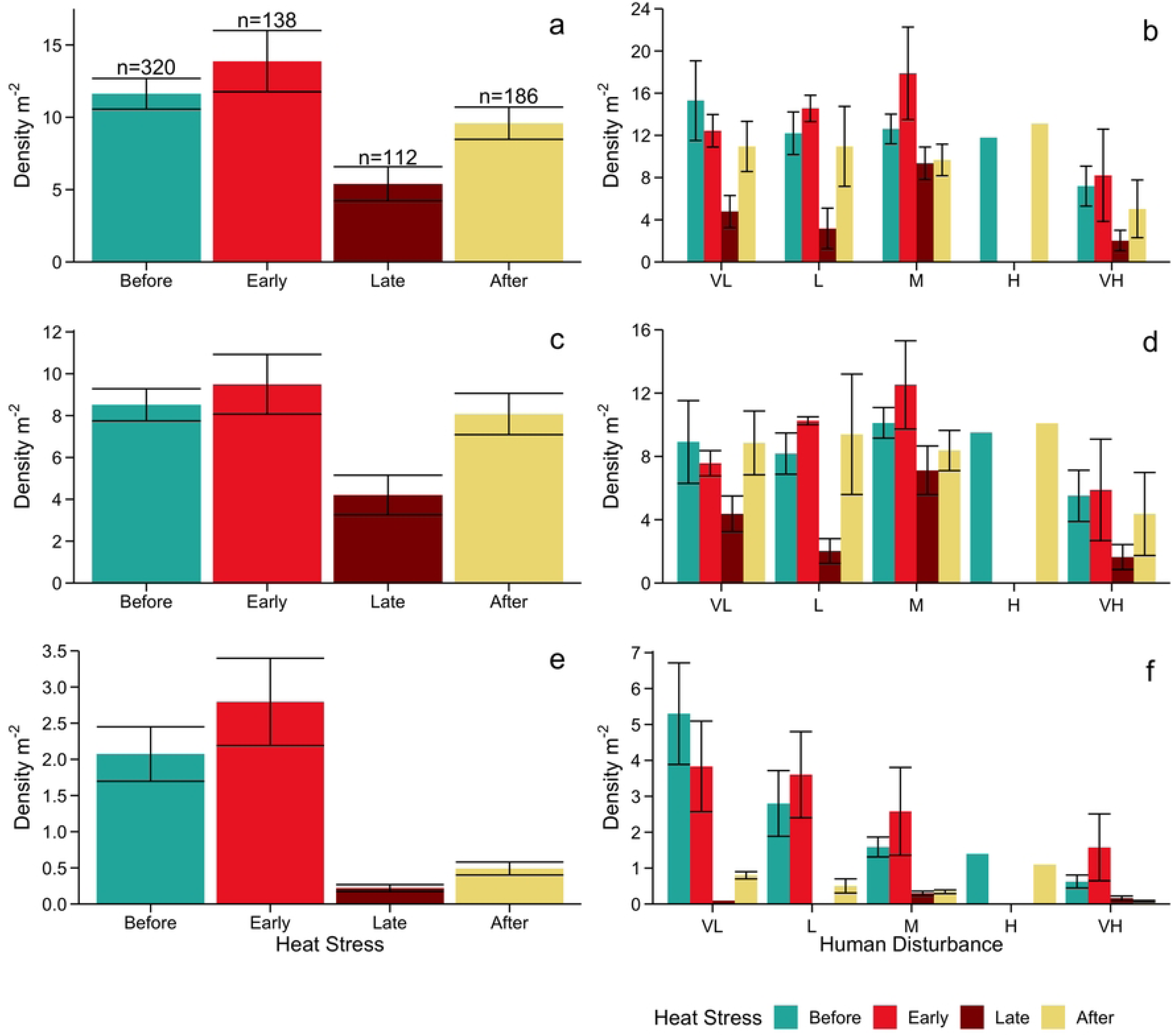
Density (± SE) of juvenile (≤5 cm) corals at forereef sites on Kiritimati (Christmas Island) across the heatwave. (Early = 2 months (July 2015), Late = 10 months (March 2016), and After = ∼1 year after (summer 2017)) for **(a, b)** all corals, **(c, d)** stress-tolerant corals, and **(e, f)** competitive corals across the **(a, c, e)** entire island (18 sites; n = the number of quadrats sampled per time point), and the **(b, d, f)** local human disturbance gradient (VL = very low, L = low, M = medium, H = high, and VH = very high). Note: Not all sites could be sampled in each time point. Y-axis scale varies among panels.

### Pre-heatwave juvenile corals

Prior to the heatwave, overall mean juvenile coral density on the atoll ranged from 7.2 m^2^ (± 1.9 SE) at the most disturbed sites up to 15.3 m^2^ (± 3.8 SE) at the least disturbed ones (Fig 4). Accordingly, pre-heatwave juvenile coral density was significantly negatively influenced by local human disturbance (parameter estimate = -0.520, *z* = -2.432, P = 0.015; S3 Table). Juvenile coral densities also varied significantly with life history strategy: stress-tolerant corals were the most common before the heatwave and declined as local human disturbance increased (S4 Fig), competitive corals also declined along the disturbance gradient (parameter estimate: -0.643, *z* = - 1.972, P = 0.049; S3 Table) but had significantly lower densities compared to stress-tolerant (parameter estimate = -1.424, *z* = -11.074, P < 0.001; S3 Table). While there were also significantly fewer weedy juvenile corals compared to stress-tolerant ones (parameter estimate = -2.499, *z* = -12.608, P < 0.001; S3 Table), they differed in that they significantly increased with increasing local human disturbance (parameter estimate = 0.642, *z =* 2.315, P = 0.021; S3 Table). Although there was a tendency for juvenile coral densities to increase with exposure (parameter estimate = 0.104, *z* = 0.462, P = 0.644; S3 Table) and NPP (parameter estimate = 0.155, *z* = 0.599, P = 0.559; S3 Table) these effects were not statistically significant.

### Heatwave effects

The 2015-2016 El Niño significantly reduced juvenile coral densities (parameter estimate = -0.774, *z* = -5.042, P < 0.0001; Figs 4a and 6, S3 Table), resulting in the loss of half of all juvenile corals by late in the event (11.6 m^2^ ± 1.1 SE before to 5.9 m^2^ ± 1.2 SE after; 50% decrease). Of the 15 coral taxa with the highest densities before the heatwave, all six with competitive life histories declined by more than 80% (three by 100%; Fig 5). There was a greater range of declines among the stress-tolerant taxa: only two, *Montipora* encrusting species and species from the Fungiidae family had losses greater than 80% and four taxa declined by less than 50%. *Dipsastraea speciosa* had the smallest loss at just over 5% (Fig 5).

**Fig 5.**
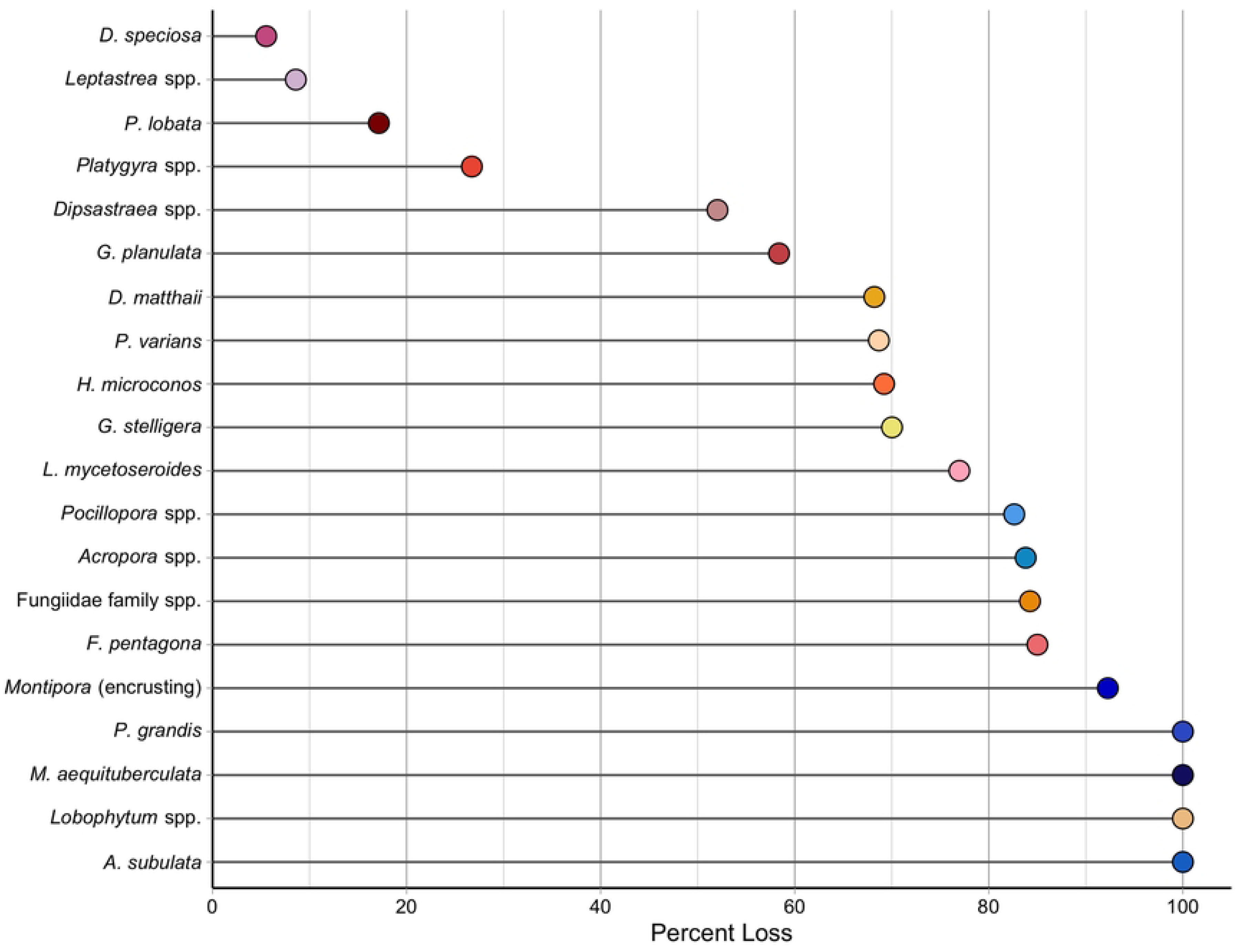
Overall change in density by the late heat stress time period of individual juvenile (≤5 cm) coral taxa at forereef sites on Kiritimati (Christmas Island) for the 15 most common taxa before the heatwave, ordered from least to greatest change in density. Where applicable, taxa colors are the same as in Baum and colleagues [31].

**Fig 6.**
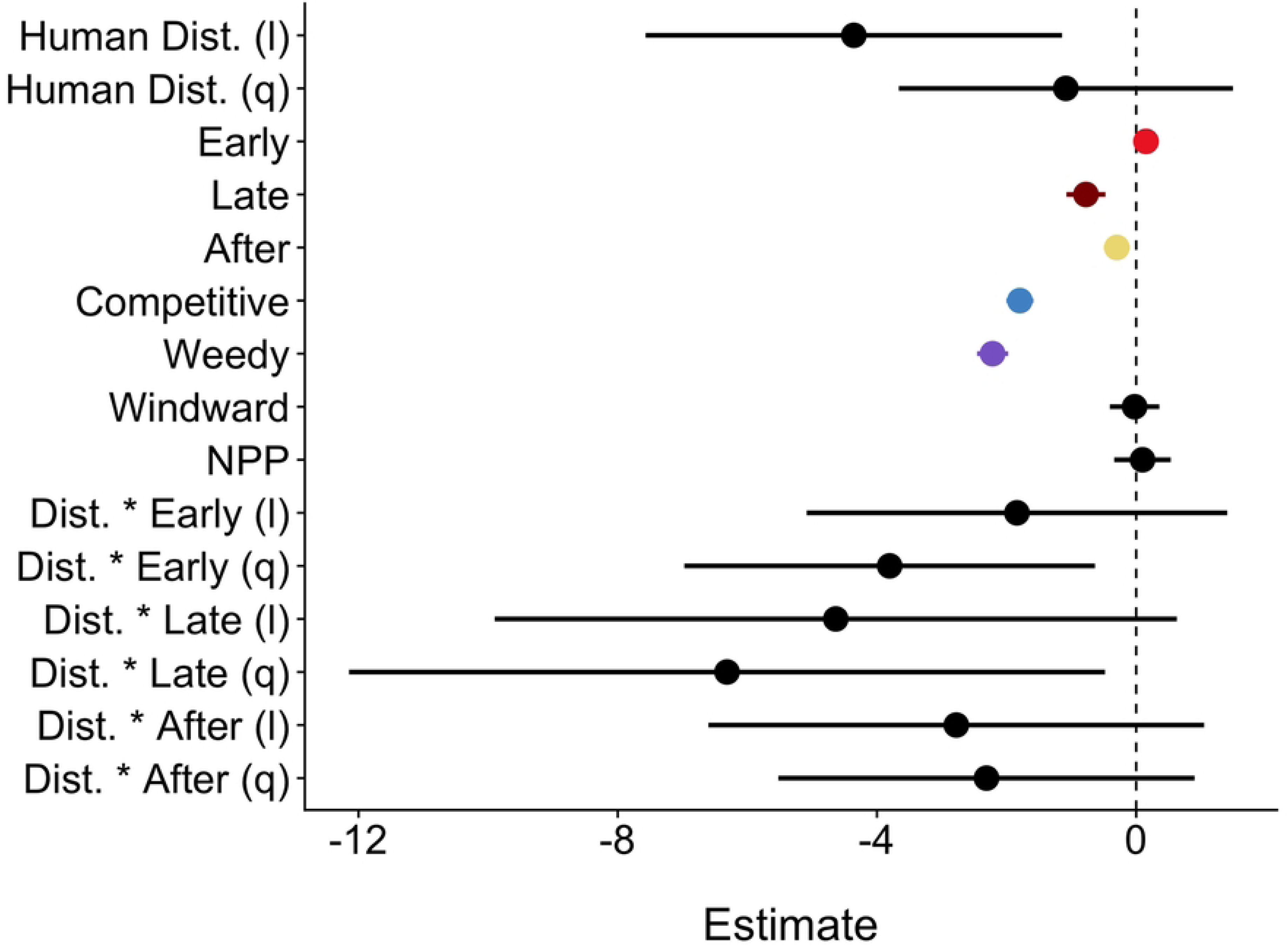
Generalized linear mixed model predictor coefficient effect size estimates and 95% confidence intervals for the overall dataset. Human disturbance was modelled as a quadratic (l= linear, q = quadratic). Heat stress colors correspond with figure 4 and life history with figure 3.

Declines in juvenile coral densities (from before to late in the El Niño) were lowest at sites in the mid-range (20.6% at medium sites, 58.5% at low sites) of the local disturbance (Fig 4). The sites at the very high and very low ends of the range experienced 66.5 per cent and 68.9 per cent respectively (high sites not sampled in late time period; Fig 4). This resulted in a significant quadratic interaction between local disturbance and heatwave for late in the heatwave (parameter estimate = -6.314, *z* = -2.121, P = 0.034; S3 Table). This quadratic interaction was also present in the juvenile coral density early in the heatwave (parameter estimate = -3.804, *z* = - 2.353, P = 0.019; S3 Table) although there were not yet decreases overall. Significant reductions in juvenile coral density were only detected late in the heatwave (March 2016; parameter estimate = -0.774, *z* = -5.042, P < 0.001; S3 Table), with none detected two months into the heatwave (parameter estimate = -0.152, *z* = 1.555, P = 0.120; S3 Table). Approximately one year after the El Niño, juvenile corals remained at significantly lower densities than prior to the event, but the effect was lessened (parameter estimate = -0.298, *z* = -2.892, P = 0.004; Figs 4a and 6, S3 Table) and densities had increased relative to those late in the heatwave (Fig 4a). The significant quadratic interaction between disturbance and heat stress dissolved by this time point (parameter estimate = -2.310, *z* = -1.409, P = 0.159; S3 Table) but there was also not a linear interaction (parameter estimate = -2.776, *z* = -1.422, P = 0.155; S3 Table). There were significantly fewer competitive and weedy juvenile corals compared to stress-tolerant corals (competitive: parameter estimate = -1.794, *z* = -16.811, P < 0.001; weedy: parameter estimate = -2.216, *z* = -18.076, P < 0.001; Fig 6, S3 Table). Neither island exposure nor NPP had significant effects on juvenile coral density (Fig 6, S3 Table). In our sensitivity analyses that controlled for site, the patterns largely remained the same with some differences in significance, notable differences include: the interaction (both linear and quadratic) between human disturbance and the late heat stress time point is significant in both site controlled models, the linear effect of human disturbance was not significant when the model was controlled for sites sampled in both before and after heat stress periods (S4 Table).

Losses in juvenile corals due to the 2015-2016 El Niño varied substantially between coral life histories: stress-tolerant juvenile corals experienced an average loss of 69.25% (parameter estimate = -0.594, *z* = -3.990, P < 0.0001; S3 Table) and weedy declined by 26.57% (parameter estimate = -0.720, *z* = -1.575, P = 0.115; S3 Table), whereas competitive coral declined by 96.8% (parameter estimate = -2.1042, *z* = -4.954, P < 0.0001; Figs 4 and S5, S3 Table). All life history strategies had an increase in juvenile corals one year after the El Niño (Figs 4 and S5, S3 Table). Across all three life history models, the results indicate that local human disturbance lowers juvenile coral density, significantly for both stress-tolerant and competitive (stress-tolerant (linear): parameter estimate = -2.487, *z* = -2.309, P = 0.021; competitive: parameter estimate = -1.169, *z* = -3.318, P = 0.001; weedy (linear): parameter estimate = -2.615, *z* = -1.874, P = 0.061; S5 Fig; S3 Table). The influence of human disturbance varied between life histories throughout the heatwave; there was no significant interaction for competitive corals suggesting that the impact of local human disturbance did not change throughout the marine heatwave. Comparatively, stress-tolerant corals had a significant quadratic relationship with human disturbance early in the heatwave (parameter estimate (quadratic) = -2.301, *z* = -2.204, P = 0.028; S3 Table) that shifted into a linear relationship by the late time period (parameter estimate (linear) = -3.088, *z* = -1.973, P = 0.049; S3 Table). Weedy corals also had a significant linear interaction late in the heat wave (parameter estimate (linear): -8.700, *z* = -1.971, P = 0.0487; S3 Table and S5 Fig, S3 Table).

Influences of island exposure varied between life history strategies. Competitive densities were significantly higher on the windward side (parameter estimate = 0.7547, *z* = 2.549, P = 0.01082; S3 Table). Weedy density patterns were the opposite with significantly less on the windward side (parameter estimate = -1.158, *z* = -3.750, P = 0.0002; S3 Table). Stress-tolerant densities patterns were slightly higher on the windward side, although not significant (parameter estimate = 0.021, *z* = 0.097, P = 0.923; S5 Fig, S3 Table).

Bleaching of juvenile corals was less than 15% at both time points measured during the heatwave (early = 14.4 ± 0.8% SE; late = 13.0 ± 3.4% SE) (Fig 7a). Before the heatwave, juvenile corals on the most disturbed reefs had the highest prevalence of bleaching (10.9 ± 1.9% SE). This shifted to reefs with medium human disturbance a couple months into the heatwave (16.2 ± 1.7% SE) and by the end, the lowest human disturbed reefs had the highest occurrence, with nearly a quarter of the juvenile corals bleached (24.8 ± 13.7% SE) (Fig 7b). For many of the most common corals surveyed, the bleaching percentage before, early, and late in the heatwave did not exceed 25%. Additionally, most juvenile coral species showed an increase in bleaching early into the heatwave but decreased by the late time period. For example, *L. mycetoseroides*, the most common juvenile coral, had 3.4% bleaching before which increased to 25.3% early on and decreased to 9.8% by the end of the heatwave (S5 Table). Seven coral species were surveyed as adults in Baum and colleagues at the same sites and time points [31]. Among those species, the juvenile corals of *Platygyra* spp. had the highest proportion of bleaching before (18.1%) and late (27.3%) in the heatwave, and second highest early in the heatwave (20%). *Dipsastraea* spp. was highest early in the heatwave (38.4%) and was second highest before (10.9%) and late (17.5%) (S5 Table).

**Fig 7.**
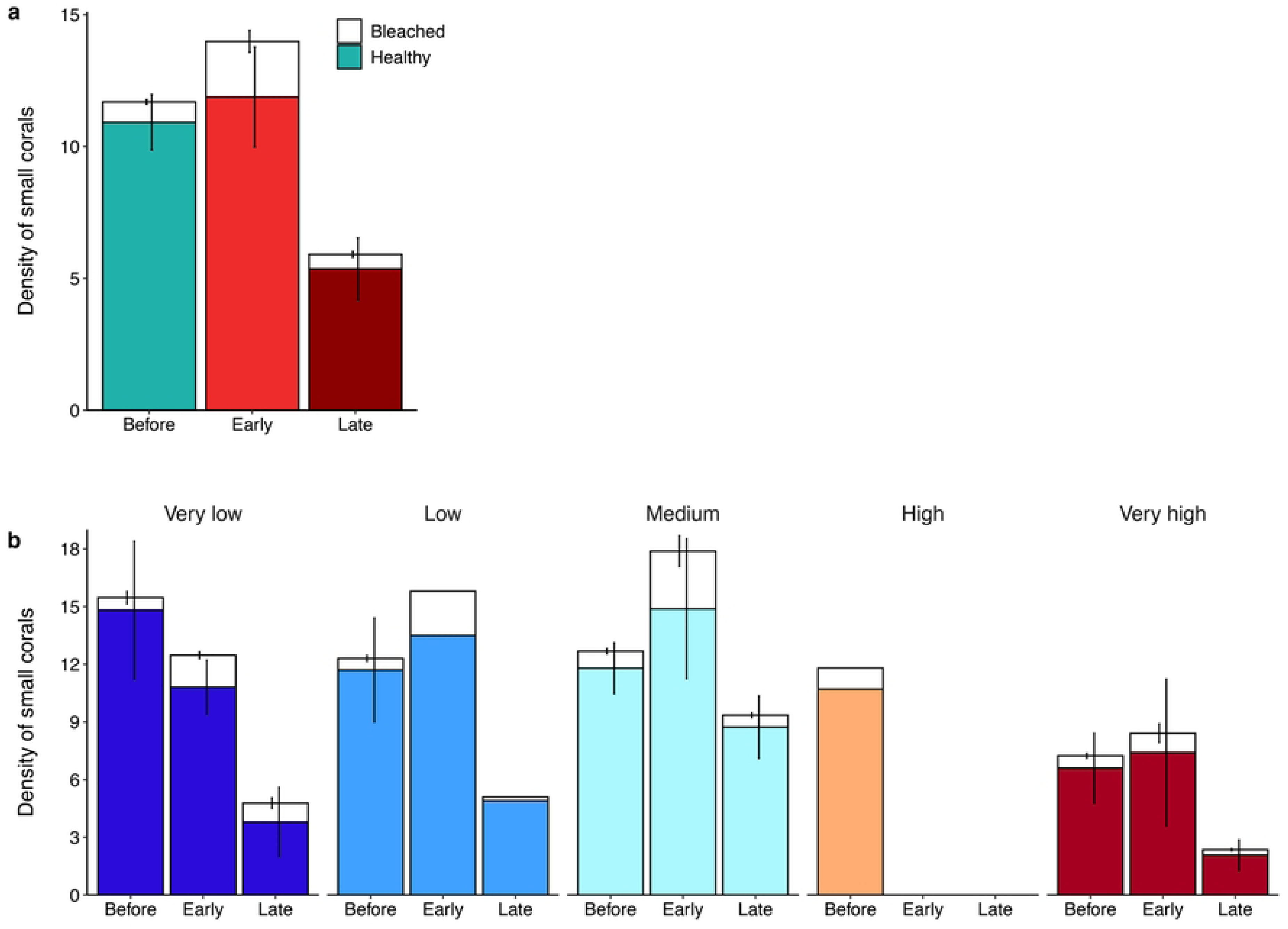
Density of juvenile corals visually bleaching or healthy across the **(a)** marine heatwave and **(b)** the human disturbance gradient. Heat stress colors correspond with figure 4 and human disturbance colors correspond with figure 1.

## Discussion

Our study provides evidence that both chronic local anthropogenic disturbance and prolonged heat stress significantly reduce the density of juvenile corals on shallow forereefs. Before the heatwave, there was an inverse relationship between juvenile coral densities and the local disturbance gradient overall, but varying impacts of disturbance on different coral life histories. The heatwave significantly reduced juvenile coral densities, and for stress-tolerant and weedy corals shifted the relationship between their density and local human disturbance. Juvenile coral densities appeared to be increasing (by >50%) just one year after the heatwave, but the differences were not statistically significant.

The significant negative effect of local human disturbance on juvenile coral densities we detected is in accordance with previous studies that have documented the negative effects single [23,27,50] and multiple [30] chronic anthropogenic stressors have on juvenile corals. On Kiritimati, this effect is believed to be the result of poor water quality, due to sewage and other pollution outflow onto the reef [31,32,51], and direct damage from dredging at some sites [31], which had substantially decreased overall coral cover and habitat complexity at the most disturbed sites prior to the heatwave [31]. We note that prior to the heatwave, turf algae and sediment cover were highest at the most disturbed sites, all of which can negatively impact corals of all life stages [21,28,29]. Comparatively, however, examining the pre-heatwave coral communities, the influence of human disturbance was greater on adults [31] than it was on juveniles (this study), although the mechanism for this difference remains unclear.

The 2015-2016 El Niño caused significant mortality of juvenile corals on Kiritimati, but this effect was only evident at our end of heatwave expedition (i.e. after 10 months of heat stress). Juvenile corals have been known to survive a few months of heat stress [15–17,52] which is hypothesized to be due to their small and flat structure increasing mass transfer off toxic by-products [19,20] and being non-reproductive [18]. Despite the apparent resilience of juvenile corals to short term heat stress, our study and others demonstrate juvenile coral’s susceptibility to long term elevated temperature conditions, similar to adult colonies [4,31,53]. Studies in the Seychelles documented 48% or greater loss of juvenile corals following the 2015-2106 El Niño [30,54] and a study in the Maldives recorded similarly low densities as our study (2.7 ± 4.6 to 5.8

± 12.3 individuals m^-2^; [13]), and although they did not have pre-heatwave data, as mentioned by Perry and Morgan [13], in the context of densities reported after the 1998 El Niño [55,56] these are very low. On Kiritimati, recovery of juvenile corals may have occurred, as evidenced by the ability of reefs on Moorea to recover despite multiple acute disturbances resulting in low juvenile coral densities [57], however subsequent monitoring has been limited due to COVID-19 related international travel bans from 2020 to 2023.

A comparison between the heatwave impacts we documented here to those documented for adult corals by Baum et al. [31] on Kiritimati, reveals that the heatwave had a greater impact on adult corals, with a higher occurrence of bleaching and 1.8 fold greater mortality in adult colonies [31]. Before the 2015-2016 El Niño, overall on Kiritimati juvenile corals were more frequently bleached than the adults [31], which could indicate that they were competing with post-settlement stressors as the escapement size, irrespective of life-history and habitat, is 5cm [7]. However, during the heatwave, the bleaching frequency in adults surpassed that of the juveniles [31], possibly due to juvenile corals’ hypothesized better resistance to bleaching [15,17–19]. In an Australian study, Álvarez-Noriega et al. [18] also found an overall difference in bleaching induced mortality between adults and juvenile corals, but that it was taxon dependent. This seems to be location dependent however, as we only found *Goniastrea* spp. to have the same trend on Kiritimati where the adults were more affected. The biggest difference was in the Merulinidae family, where in Australia the juveniles did worse than their adult counterparts, but on Kiritimati, the majority of the Merulinidae juveniles increased in density while the adults declined [31]. Small colonies of *Oculina patagonica* also had higher survivorship than their adult counterparts during a bleaching event in the Mediterranean [52], whereas in the inner Seychelles, Dajka and colleagues [30] documented 70% mortality of juveniles due to heat stress which was similar to the adult community loss on those reefs.

It was expected that the corals at the sites exposed to very high local disturbance before the heatwave would have been the most stressed and therefore have the highest prevalence of bleaching, however, the greatest prevalence of bleaching shifted along the local human disturbance gradient throughout the heatwave. Interestingly, as the shift in high bleaching prevalence moved to medium disturbed sites early in the event, it did not correspond with a large die off of juvenile corals at the very high disturbed sites. In fact, there was an increase in juvenile corals between the before and early time points. Additionally, while there was a large decline in juvenile corals by the late time period across the atoll, the sites that experienced the highest bleaching percentage (medium disturbed sites) at the early time point had the smallest decline (∼21%). This demonstrates a mismatch between bleaching prevalence and mortality levels similar as to what was recorded in the adult colonies [31]. Mechanisms for this have been proposed for adult colonies (e.g., resistant to bleaching but then can only survive temporarily in a bleached state [58,59], symbiont switching mid heat event [35]) but to our knowledge this pattern has not been documented in juvenile corals. It is possible that juveniles follow some of the same mechanisms as adults but with their different physical structure [19], they may have other mechanisms.

Juvenile corals with a stress-tolerant life history were dominant across the atoll at all time points in this study. While for the entire coral community, Baum et al. [31] also found that stress-tolerant corals dominated Kiritimati’s highly disturbed reefs and all reefs after the 2015-2016 El Niño, prior to the event, competitive corals dominated the lowest disturbance sites. Dominance of stress-tolerant corals was also documented on reefs in the Maldives in the years immediately following both the 1998 [55] and 2016 El Niños [13]. In contrast, after heatwaves other reefs were dominated by competitive and weedy type corals (e.g., *Acropora* and *Pocillopora*) that colonize open space on reefs and grow rapidly [10,16,57,60]; however, the timescale varies globally, likely due to differences in community dynamics and severity of disturbances. Our study only extended one year past the El Niño and we detected no significant increase in weedy and competitive corals compared to stress-tolerant corals, similar to what was recorded on nearby Jarvis Island [61]. This might indicate that recovery will be suppressed until these corals can contribute significantly to recruitment, as documented on other reefs severely impacted by heat stress [7,10,12,13,62] compared to reefs less impacted by extreme heatwaves [57,63]. Thus, with more studies, it may be possible to roughly calculate recovery times using the severity of the disturbance and composition and density of surviving juvenile corals.

Reef exposure has been shown to protect juvenile coral densities during marine heatwaves as water velocity is a rate-determining step in mass transfer [20] however, high wave energy can have a negative impact during recovery [64–66]. In this study, as on the Great Barrier Reef [7], habitat specific stressors varyingly influenced juvenile assemblages. The sites located along the windward shores had higher densities of competitive corals possibly due to these sites being dominated by adult competitive corals [31] and coinciding with a lack of local human disturbance resulting in low competition with macroalgae. In contrast, weedy corals had higher densities on the leeward and more disturbed shores, which is a common trend around the world [7,24,67–70].

Overall, our study demonstrates the negative impacts that a prolonged heatwave and localized chronic anthropogenic stress had on juvenile corals. This study also highlights differences in impacts amongst juvenile corals with different life history strategies. Given the importance of juvenile corals for reef recovery, continued studies of the impacts of these stressors, the mechanisms by which they affect juvenile corals, and ongoing monitoring of their contributions to reef recovery trajectories is needed.

## Acknowledgements

Thanks to Kiritimati Field Teams for collection of GoPro videos; to Roxanne Le-Goff for assistance with video analysis, Jacob and Lavina Teem and the crew at Ikari House for taking care of us on island, the Kiribati Government for their support of this research, and to the Kiribati people. We also acknowledge and respect the Lək̓ʷəŋən (Songhees and Esquimalt) Peoples on whose territory the University of Victoria stands and the majority of the authors live, and the Lək̓ʷəŋən and W̱SÁNEĆ Peoples whose historical relationships with the land and waters continue to this day. Research was conducted under research permits: 008/13, 007/14, 001/16, 003/17.

## Supporting information

**S1 Fig. Map of reef study sites on Kiritimati (Christmas Island) sampled at each heat stress time point** (Before = 2 years before – start (summer 2013, summer 2014, April/May 2015), Early = 2 months (July 2015), Late = 10 months (March 2016), and After = ∼1 year after (summer 2017). The sites are divided into five levels of local human disturbance. Village population (red circles) is represented by bubble size.

**S2 Fig. Representative photos of quadrats surveyed on Kiritimati (a, d)** before, **(b, e)** at the end, and **(c, f)** after the heatwave on reefs exposed to **(a-c)** very high levels of local human disturbance and **(d-f)** very low levels.

**S3 Fig. Widest width of juvenile corals (n=7732) on Kiritimati throughout the survey.** Mean width was 2.28 cm (± 0.01 SE). Bins are 0.5cm.

**S4 Fig. Density of juvenile corals with different life history strategies across the human disturbance gradient prior to the 2015-2016 marine heatwave** modelled with a two-way interaction.

**S5 Fig. Generalized linear mixed model predictor coefficient effect size estimates and 95% confidence intervals** for each tested life history strategy: **(a)** stress-tolerant, **(b)** competitive, and **(c)** weedy. In models a and c, human disturbance was modelled as a quadratic (l = linear, q = quadratic). Heat stress colors correspond with figure 4. X-axis scale varies among panels.

**S1 Table. Number of juvenile coral video assays conducted at each of 18 sites around Kiritimati, by expedition before (July 2013, August 2014), during (May 2015, July 2015, March 2016), and after (July 2017) the 2015-2016 El Niño.** Sites are ordered first by decreasing levels of local chronic human disturbance then by exposure (Fig 1).

**S2 Table. Life history table of juvenile coral taxa identified from video assays processed using Tracker.** Coral life history strategy retrieved from the Coral Traits Database (https://coraltraits.org/), unless otherwise noted. Current taxonomy (and name synonymy) retrieved from WoRMS (http://www.marinespecies.org/).

**S3 Table. Results for (a) pre-heatwave and (b) heatwave models.** Bolded values are significantly different from baseline levels (i.e., before, stress-tolerant, leeward) at α = 0.05, asterisks indicate levels of significance (· *p* < 0.1, * *p* < 0.05, ** *p* < 0.01, *** *p* < 0.001). Red shaded boxes correspond to variables with a negative parameter estimate.

**S4 Table. Results for the sensitivity analysis on the heatwave effect by controlling for sites sampled.** The first model **(a)** was run using only the 9 sites that were sampled in all four heat stress periods (S1 Fig). The second model **(b)** is run with the 10 sites that were sampled in both the before and late heat stress periods. Bolded values are significantly different from baseline levels (i.e., before, stress-tolerant, leeward) at α = 0.05, asterisks indicate levels of significance (· *p* < 0.1, * *p* < 0.05, ** *p* < 0.01, *** *p* < 0.001). Red shaded boxes correspond to variables with a negative parameter estimate.

**S5 Table. Numbers of each species identified in the video assays during each time point and overall.** Current taxonomy retrieved from WoRMS (http://www.marinespecies.org/).

**S5 Table. Percent bleaching for juvenile (JV) corals during three time points for top 16 coral species and two other species** of which there is adult data to compare to. Rank column denotes level of common-ness before the heatwave; the 7^th^ most common was unidentified and thus not included on this table. Adult data was only available for seven species from Baum and colleagues [31]. (n = total for that time period)

